# Seeing Double: Molecular dynamics simulations reveal the stability of certain alternate protein conformations in crystal structures

**DOI:** 10.1101/2024.08.31.610605

**Authors:** Aviv A. Rosenberg, Sanketh Vedula, Alex M. Bronstein, Ailie Marx

## Abstract

Proteins jiggle around, adopting ensembles of interchanging conformations. Here we show through a large-scale analysis of the Protein Data Bank and using molecular dynamics simulations, that segments of protein chains can also commonly adopt dual, transiently stable conformations which is not explained by direct interactions. Our analysis highlights how alternate conformations can be maintained as non-interchanging, separated states intrinsic to the protein chain, namely through steric barriers or the adoption of transient secondary structure elements. We further demonstrate that despite the commonality of the phenomenon, current structural ensemble prediction methods fail to capture these bimodal distributions of conformations.

## Introduction

Despite the dynamic nature of protein molecules, X-ray crystallography, perhaps the most widely used method for studying their structure, produces only static structure models. The ensemble nature of this measurement modality does, however, provide some information about the structure flexibility through the B-factors and electron density, and these structures allow specifying *alternate locations* for atoms to represent the dynamics (Stachowski and Fischer, 2023, Djinovic-Carugo 2015, Gutermuth 2023). For example, side chains are commonly found and modelled as a population of a discrete set of rotamer conformations, (Figure 1). Multiple slightly altered backbone conformations observed in high resolution structures have been associated with these alternate side chain rotameric states (so-called backbone shrugging, Davis et al. 2006). Other distinct alternate backbone conformations have been observed and attributed to the crystal environment (Sevick et al. 2004, Tanabe et al. 2020; Figure 1), or interactions with ligands or other binding partners. When ligands are only bound to a portion of the protein molecules (partial occupation), the crystal electron density may appear as a mixture of bound and unbound states and thus modelled as two alternative conformations of the backbone (Schlieben et al. 2005). Similarly, proteins have crystallized from a mixture of oxidized and reduced forms, and the sections of the backbone harboring the cysteines that form a disulfide bond are modelled in alternate conformations (Pernigo et al. 2010; Figure 1).

**Figure 1.**
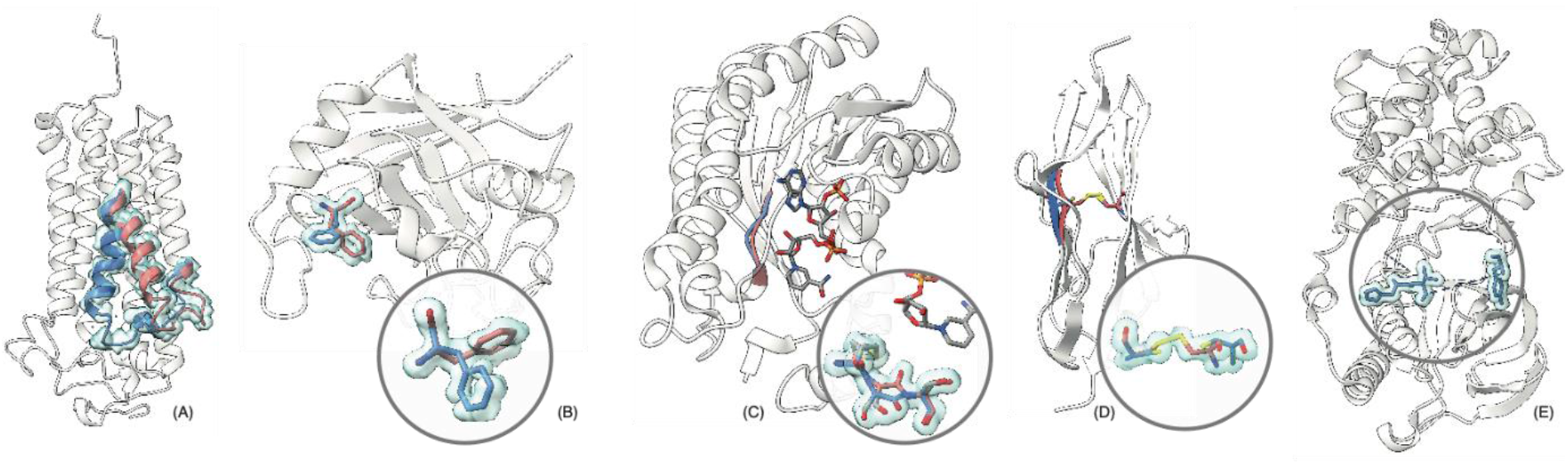
Examples of alternate conformations with environmental explanations. (A) a protein in two states (open and closed) mediated by allosteric effects; (B) a population of two discrete rotamer states of a sidechain; (C) bound and unbound states due to ligand interaction; (D) different oxidation states of a cysteine pair. Furthermore, some regions in many protein structures exhibit such a degree of flexibility, that no accurate modelling is possible (dashed lines, E). The backbone conformation cannot be determined with crystallographic methods in such cases.

In this work, we are interested in alternate conformations that do not fall into the above categories. We apply a large-scale analysis to identify well-separated alternate local backbone conformations, modelled in single crystal structures, that cannot be trivially explained by proximity to ligands, binding partners, or crystal contacts. We demonstrate, through molecular dynamics simulations, that well-separated alternate conformations often remain stable, thus effectively creating distinct molecules. Long-lived metastable structures, kinetically trapped in the protein folding funnel, have previously been detected by atomic force microscopy (Shin et al. 2012), NMR (Gautier et al. 2020), and crystallography (Hua et al. 1995).Our results suggest that these phenomena may be more common than previously realized.

According to the OCA browser database for protein structure/function (Prilusky, 1996), over 95% of Protein Data Bank (PDB) structures deposited after 2010 contain backbone atoms with alternate locations (altlocs). Gutermuth (2023) recently noted that the prevalence of altlocs is often unnoticed since most modelling approaches either ignore altlocs altogether or resolve them with simple heuristics. This could bear substantial unintended consequences, for example when such data are used to train structure prediction models such as AlphaFold (Jumper et al. 2021) as we shall discuss below. We recently compiled a comprehensive and detailed catalog of alternate backbone segments from the entire PDB, aiming to bring attention to this matter and facilitate proper analysis of structures with altlocs (Rosenberg et al. 2024). Figure 2 demonstrates the abundance of altlocs in PDB structures. Based on the highest resolution structures, we find that over 4% of proteins harbor non-terminal backbone altlocs exhibiting a multi-modal electron density distribution (Supplementary Figure 1) that cannot be explained by other factors, being are at least 4Å away from ligands, out-of-chain binding partners, or any crystal lattice contacts (Supplementary Figure 2).

**Figure 2.**
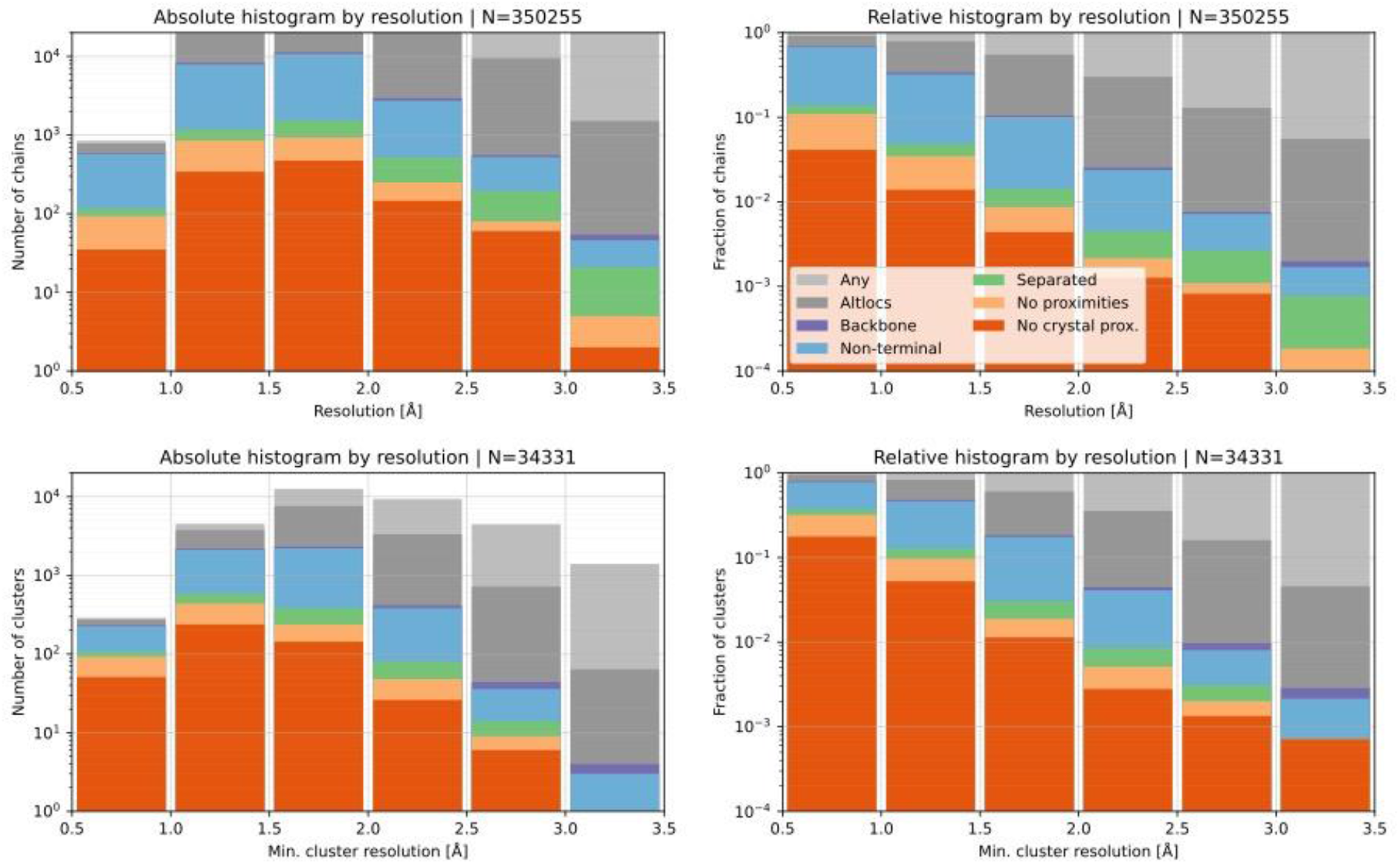
Categorisation of altlocs in over 350K crystallographic structures in the PDB. Shown are absolute and relative distributions of individual chains (first row), and non-redundant clusters (second row) broken down by resolution. Sample sizes (N) are indicated for each plot. Note that categories and the bars representing them are nested (for example, the “Separated” bars are fully included in the “Non-terminal” bars, etc.) Dark orange bins suggest a, perhaps, surprisingly high number of “one sequence, multiple structures” cases that are not readily explained by interactions or environmental factors.

Molecular dynamics (MD) has proven a powerful tool for simulating the conformations adopted by proteins fluctuating within a thermodynamic energy well surrounding a stable minimum. We employed MD to investigate the distributions of these alternate backbone conformations and quantify the intrinsic stability of the observed backbone twists within the protein chain. From the 1109 structures making up the innermost nested group in Figure 2 (dark orange), i.e., those harboring separated backbone altlocs that are non-terminal and non-proximal to ligands, out of chain contacts or crystal contacts, we selected a non-redundant subset suitable for molecular dynamics: protein chains less than 300 amino acids in length and having no breaks. To avoid non-proximal allosteric binding effects we discarded structures with any ligands outside a list of common crystallization reagents (see the Methods section). These selection criteria resulted in a list of 48 structures (see Supplementary Table), on which we ran pairs of short (50ns) MD simulations, one starting from a structure with the first altloc conformation (denoted as A and visualized in blue), and the other with the second altloc conformation (denoted as B and visualized in red). We categorized the simulation outcomes into four categories exemplified and visualized in Figure 3 (the full set is available in the Supplementary Materials).

**Figure 3.**
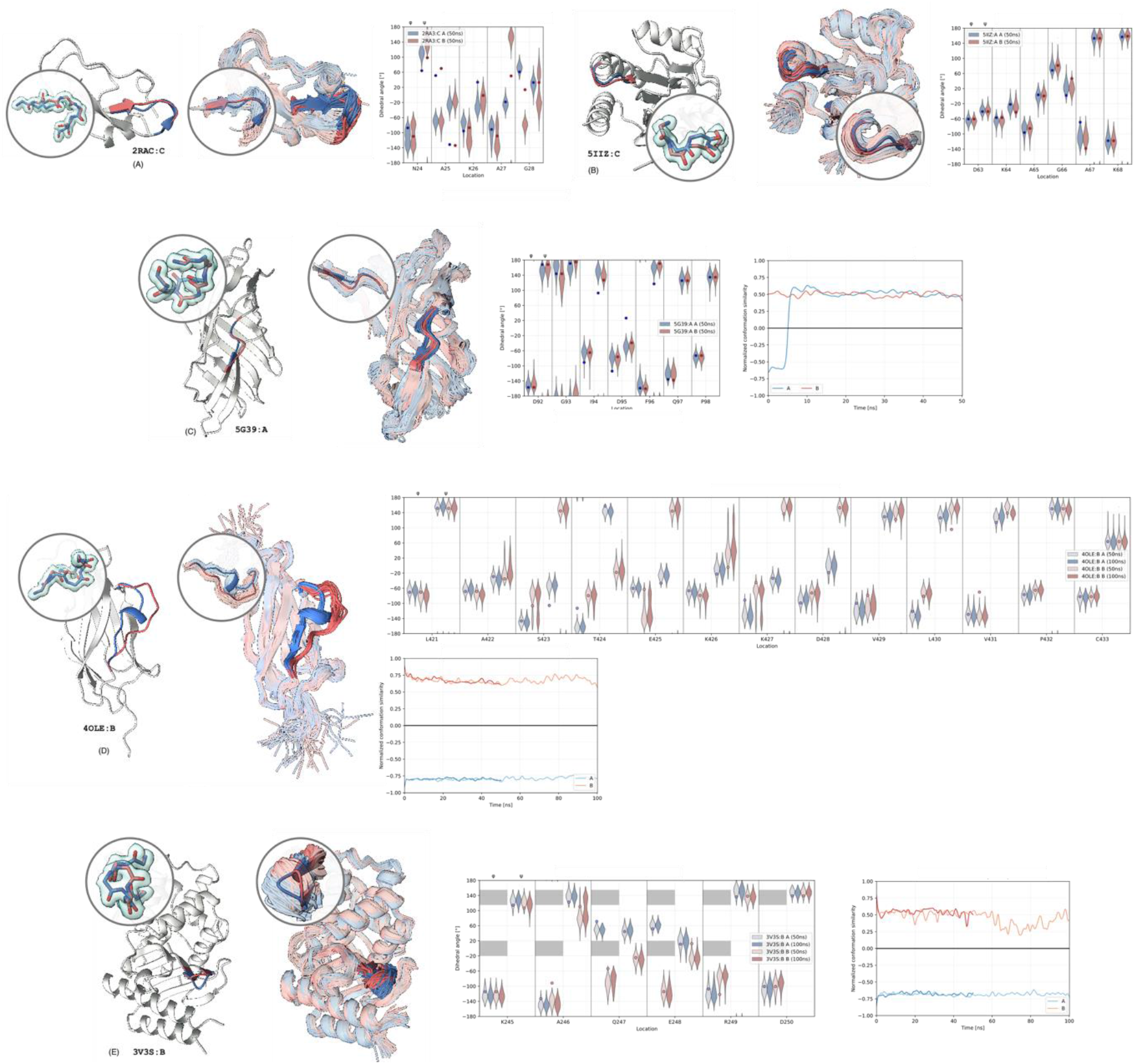
Examples of MD simulation outcomes. (A) *Failed* simulations, where both initializations converged to a third conformation that is neither A nor B; (B) *Inseparable* simulations, where both initializations produced indistinguishable ensembles; (C) *Unstable* simulations, in which one of the initializations appeared unstable and converged to the other conformation in either 50 or 100 ns run; (D)-(E) *Stable* simulations, in which both initializations produced stable and distinct ensembles resembling the appearance of the electron density. Structures in (D) are distinguished by the presence of a stable secondary structure (a helical loop in the visualized example) while structures in (E) are distinguished by a steric hindrance (visualized as shaded regions in the violin plot). Each panel displays left-to-right the structure with the two backbone altlocs segments highlighted in blue (conformation A) and red (conformation B), with a magnification showing how they fit the electron density (1σ iso-surface on 2Fo-Fc displayed in transparent cyan), an ensemble of conformations sampled from the two MD simulation runs initialized at the two conformations (with a magnification showing an overlay of the ensembles with the conformations modelled in the PDB), and the distribution of the backbone dihedral angles. Panels (C)-(E) further show the temporal trajectory of the conformation similarity (−1 corresponding to conformation A, +1 to conformation B, and 0 meaning equidistant from A and B). The violin plots show the dihedral angle distribution for all residues in the altloc segment during the simulation. Shaded regions correspond to the disallowed angles in a Ramachandran plot.

First, we observe a subset of structures that adopt conformations distinct from either the A or B starting conformations; we consider these to be *failed* cases. This is not surprising since our process can reasonably result in structural re-equilibration: structures are introduced into our standardized pipeline after being detached from their environment; isolated from all other out-of-chain entities either in the unit cell or in the crystal lattice and introduced into a standard solvent environment. Eleven structures that converged to conformations distinct from the A and B starting points were excluded from further analysis. Divergence of glycine and glycine-adjacent residues from the starting conformation was allowed since glycine is generally conformationally flexible. The second outcome was that in fourteen cases where the distributions of the A and B conformation overlapped and became *inseparable*. Although we selected well-separated altlocs using a normalized distance that accounts for the position uncertainty implicit in the B factor (see Methods and Supplementary Figure 1), these structures nonetheless became indistinguishable during simulation, which could be attributed to increased thermodynamic fluctuations in the isolated chain as compared to the crystal environment. The remaining 23 structures could be split into either *unstable* (10 structures) or *stable* (13 structures) outcomes. In the unstable cases, either the A or B conformation reverted from the starting conformation to eventually adopt the alternate conformation. Finally, in the stable cases, the initial A and B conformations were both maintained, resulting in non-overlapping distributions during the entire simulation. Structures that remained stable in the short MD simulations were simulated for an additional 100ns using different random seeds. Two structures demonstrated instability in this second, longer, simulation and a third structure adopted conformations distinct from either the A or B starting point. Figure 3 displays an example of each of the different outcomes of MD and Supplementary files 1–4 show the results for all the analyzed structures.

Analysis of the stable structures (summarized in Table 1) revealed a common feature: at least one residue in each altloc segment had φ backbone dihedral angles (for the A and B conformations) that were separated by disallowed regions in the Ramachandran plot (shaded in grey on the violin plot of Figure 3E). The prevalence of these φ*-separated* stable altlocs suggests that the alternate conformations may remain stable due to steric restrictions. Ten out of thirteen stable structures exhibited φ-separation compared to four out of ten structures in the unstable group. Of these four, one changed conformation during the longer 100ns simulation and another two had their φ-separated alternate conformations at a glycine residue. Glycine residues, having no side chain, are less sterically restricted and occupy larger regions of the Ramachandran plot. They would therefore be expected to traverse the φ-separated regions more easily. To test whether the long-lived stability of dual conformations is indeed facilitated by steric hindrance preventing exchange between the conformations, we mutated each stable structure to glycine at the point exhibiting φ-separation and ran MD simulations on the mutants (Figure 4A). As expected, in all cases the glycine mutants became unstable, with one conformation reverting to the other.

**Table 1.**
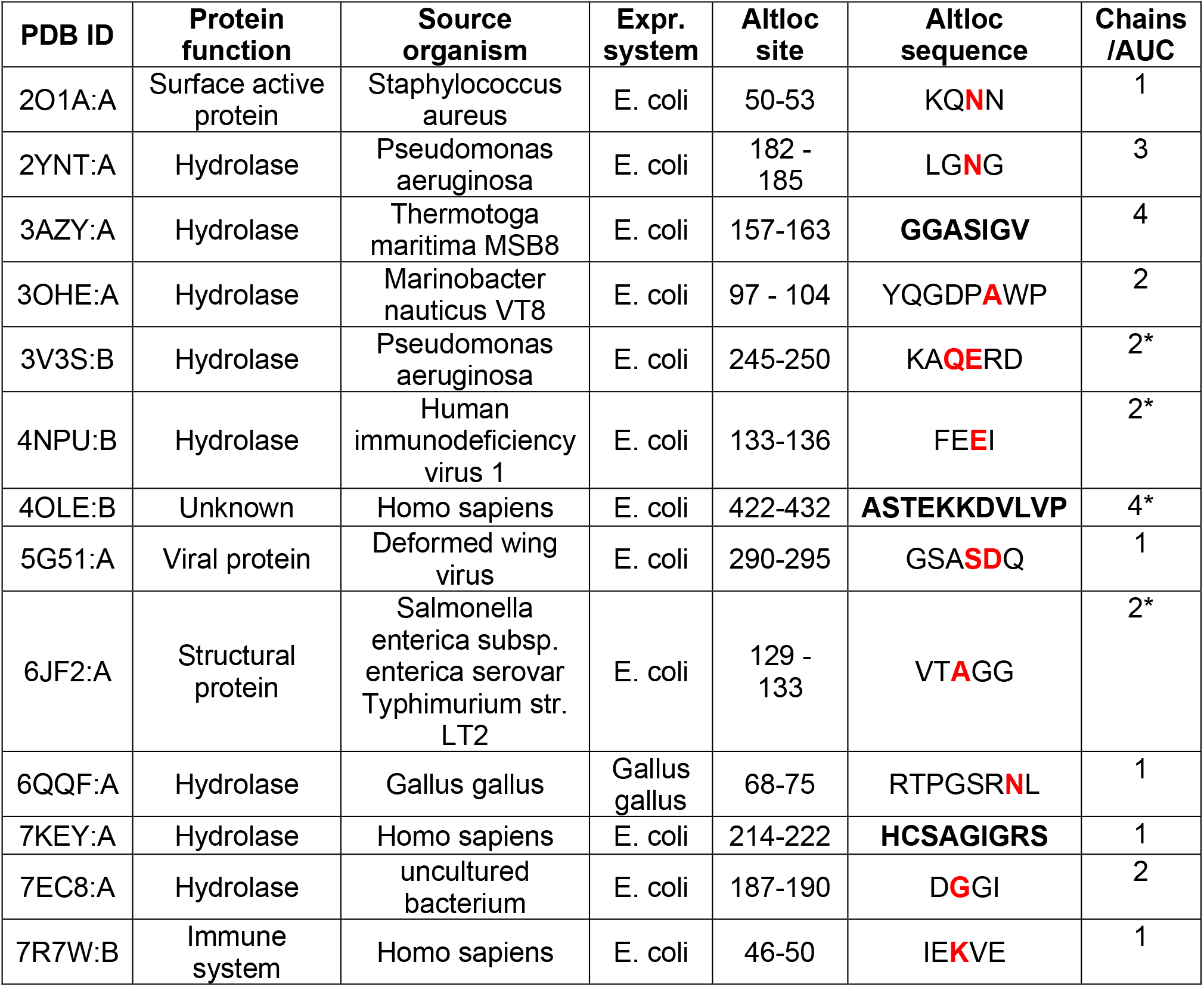
Proteins harboring stable dual backbone conformations. The alternately located segment position and sequence are indicated. Marked in red are residues modeled with φ-separation, i.e., angles on opposite sides of allowed regions in the Ramachandran plot. Bolded are sequences forming secondary structures in only one alternate conformation. The number of same chains in the asymmetric unit cell (AUC) are shown and where multiple chain exist a star indicates that the altloc is consistently present across all chains.

**Figure 4.**
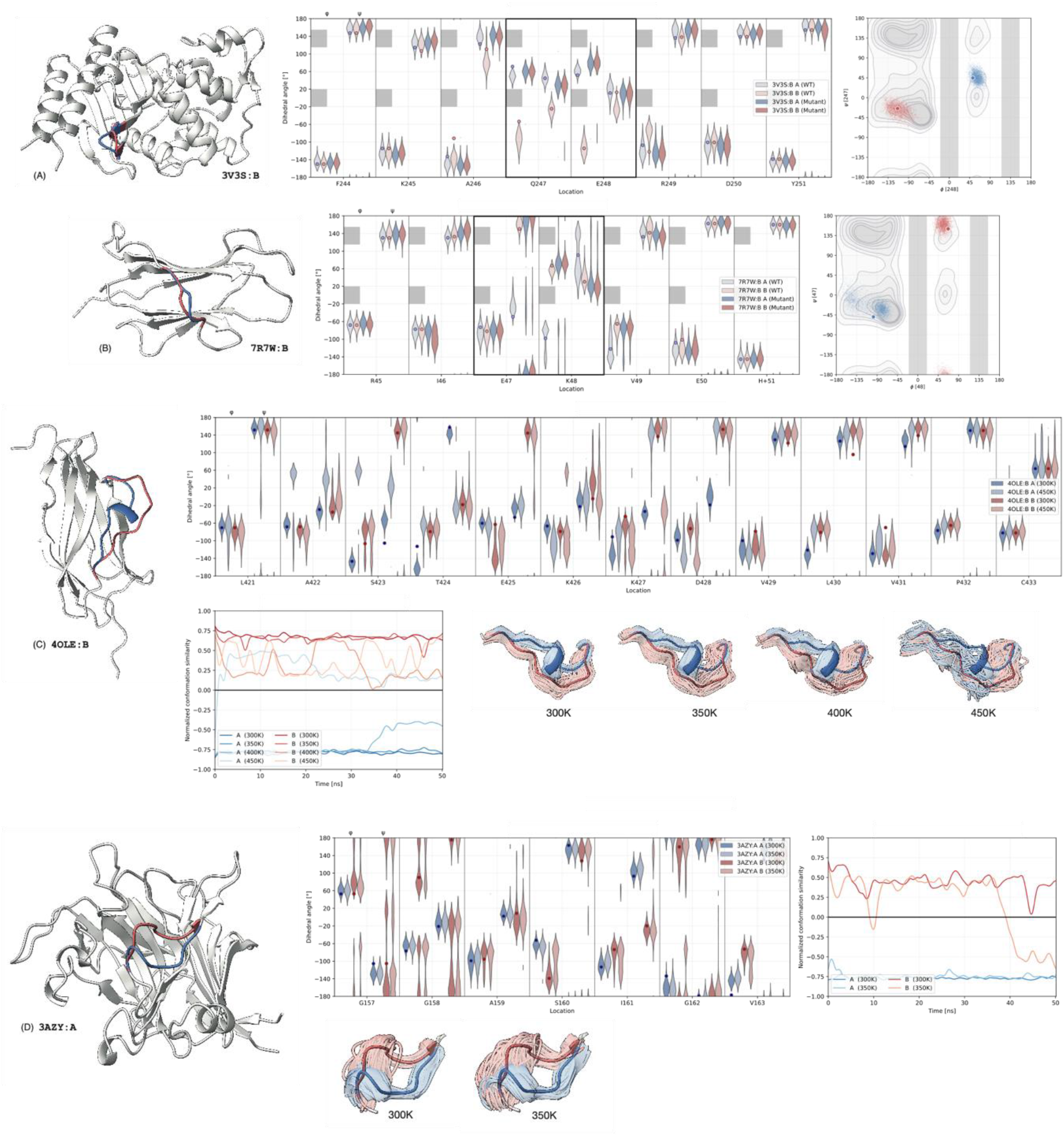
Analysis of stable dual conformations. Panels (A)-(B) show two cases where conformation stability is attributed to steric hindrances, visualized as shaded regions in the angle distribution plots and the Ramachandran plots. Mutation of the highlighted pair of amino acids to GG removes the hindrance and, as a result, the bi-modal behavior disappears. Each panel displays, left to right: the structure with the two backbone altlocs highlighted in blue (conformation A) and red (conformation B); the dihedral angle marginal distributions in 50 ns MD simulations of the wild type and the mutant initialized at each of the two conformations; the cross-bond Ramachandran plot of the wild type in two initializations. Note that the crossbond Ramachandran was employed as a better visualization of the steric restrictions, following Rosenberg *et al*. (2023). Panels (C)-(D) show two cases where conformation stability is attributed to the presence of a stable secondary structure in one of the alternative conformations (a helix in the first case and a strand in the second one). MD simulation of the structures at higher temperatures breaks the secondary structure and allows exchange between the two modes. Each panel shows, as before, the protein structure with the dihedral angle distributions, the temporal evolution of the conformation similarity at different temperatures (−1 corresponding to conformation A, +1 to conformation B, and 0 meaning equidistant from A and B), as well as structural visualizations of the simulated ensembles, overlaid on the two modelled altlocs.

The three remaining stable structures, which did not exhibit φ-separation, all revealed a second feature that potentially stabilized their dual conformations: intact vs. frayed secondary structure. Figure 3 (D) shows a clear example: the A conformation of 4OLE adopts an α-helical turn, while the B conformation of the corresponding segment appears as an open loop. The A conformation thus benefits from the stabilizing hydrogen bonds that are part of the secondary structure it forms, while the B conformation does not. To explore the stabilizing effect of the hydrogen bonding within the altloc segments, we simulated these three structures at increasing temperatures in standard molecular dynamics. We found that the added energy caused the α-helical turns to open, after which the segments adopted a conformation similar to the B altloc. Incremental heating captured a partially open state, suggesting that the transition between the A and B conformations happens as a sequential loss of the α-helical stabilizing hydrogen bonds (Figure 4B).

The structures harboring stable dual alternate conformations are listed in Table 1. Of note is that these structures appear in different protein families derived from a wide variety of species. In addition, the crystal structure 6QQF, natively expressed Hen white lysozyme, was measured at room temperature. Observing stable, long-lived alternate conformations in a room temperature structure of such an intensely studied protein might support the suggestion that cryo-cooling of protein crystals masks conformational ensembles that would otherwise be observable in room temperature experiments (Keedy et al 2014, Fraser et al 2011). It is also worthwhile noting that half the structures having multiple chains in the asymmetric unit cell display consistent dual conformations across the chains, further suggesting the latter being an intrinsic property of the chain rather than an environment-induced phenomenon (e.g. 4OLE, Supplementary Figure 3).

Computational structural biology is actively looking beyond the “one sequence – one structure” dogma with numerous algorithms now producing an ensemble of protein structures given an input sequence. We evaluated the ability of Alphafold (Jumper et al 2021 and Abramson et al 2024) and recent conformation sampling algorithms (Jing et al., 2024 and Zheng et al 2024), to capture the stable alternate conformations identified in this work. Figure 5 shows that although the global structure of the proteins was predicted with high accuracy, none of the algorithms were successful in predicting the bi-modal behavior of the backbone in the altloc segments of interest, either modeling these locations as largely flexible or capturing only one of the two modes.

**Figure 5.**
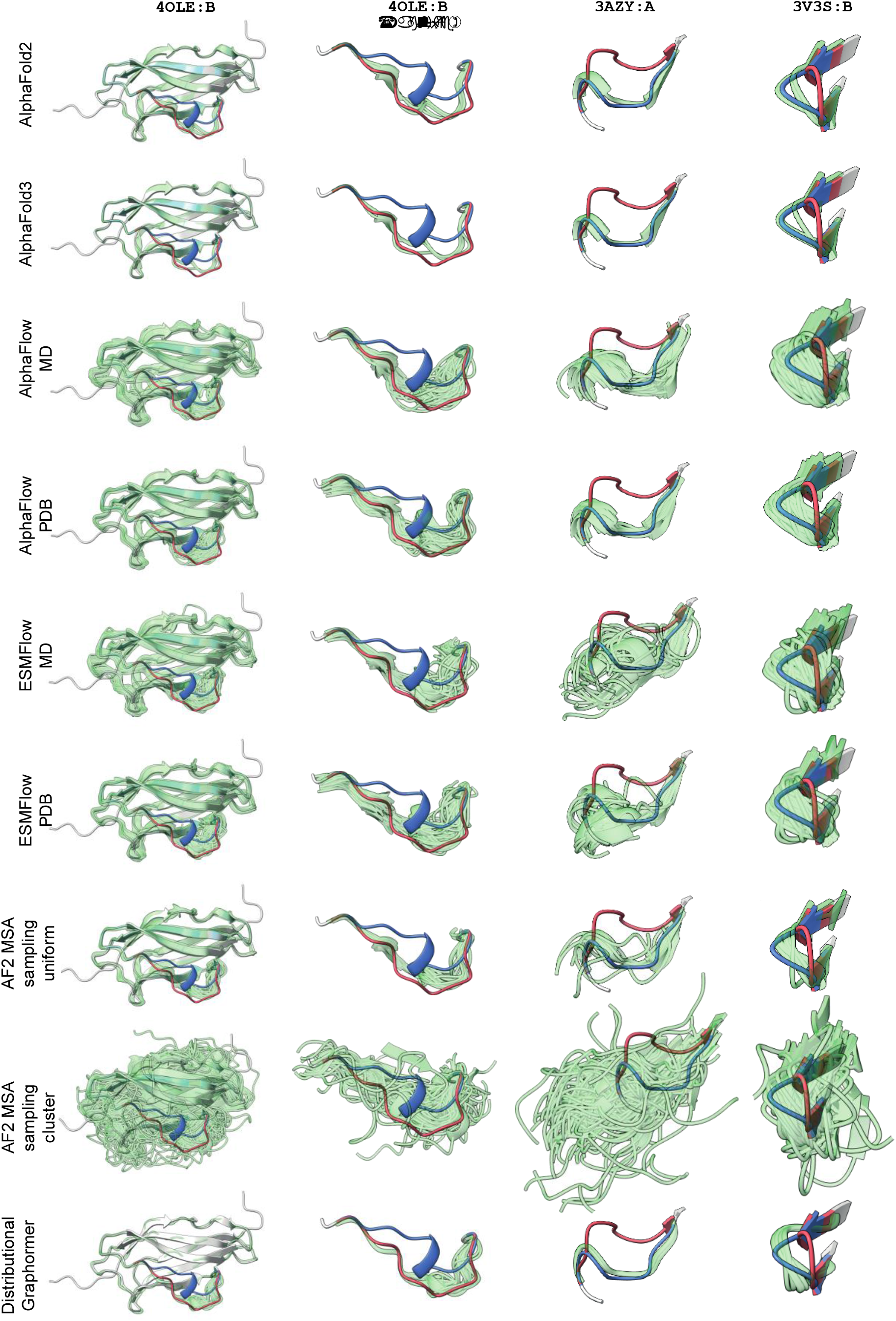
Failure of ensemble prediction algorithms in altloc segments. Although structure and ensemble prediction techniques based on amino acid sequences are capable of accurately reconstructing the overall backbone structure (left column), they consistently fail to produce viable samples corresponding to one of the two altloc conformations (as shown in the magnified regions in the second through fourth columns).

In conclusion, we have identified a set of structures that demonstrate dual stable conformations, a clear example of different conformations arising from the same amino acid sequence. These structures also provide concrete targets for experimentally probing into protein folding and the formation of metastable states. Moreover, we believe these structures provide an excellent benchmark for ensemble structure prediction methods, which aim to produce faithful samples from the conformation space given an input sequence. We note that the identified proteins were the product of a conservative pipeline aimed at removing any potential confounding factors. Revisiting this analysis and including sites close to crystal or out-of-chain contacts would likely reveal many more cases that remain stable in isolation. As is the case for any other region in the protein chain, it is often the protein conformation that facilitates the interaction and not the interaction that creates the conformation. In this vein, it is also interesting to explore dual stable conformations in proximity to ligands and ligand binding sites. Recently, MD simulations were used to show how mutations, known to affect activity, can affect the distribution of alternately located side-chain conformations (Palaniappan et al 2024). Identifying dual conformations in the backbone of ligand binding sites could lead to new insights into the mechanisms that modulate protein activity.

## Acknowledgements

AM acknowledges the financial support of the Helmsley Fellowships Program for Sustainability and Health. AMB is supported by the Schmidt Chair in Artificial Intelligence.

## Methods

### Altloc segment categories

We categorized contiguous altloc segments into the following 6 nested categories:

Crystal contacts were calculated using Pymol (Schrödinger and DeLano 2020) symmetry operators within a 4Å threshold.

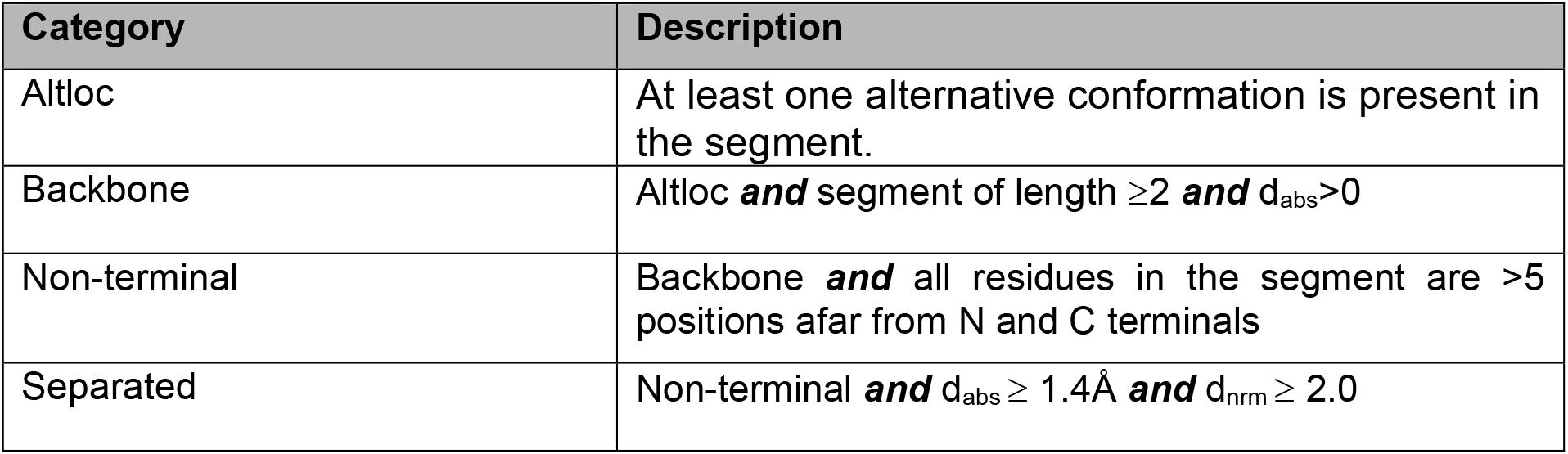
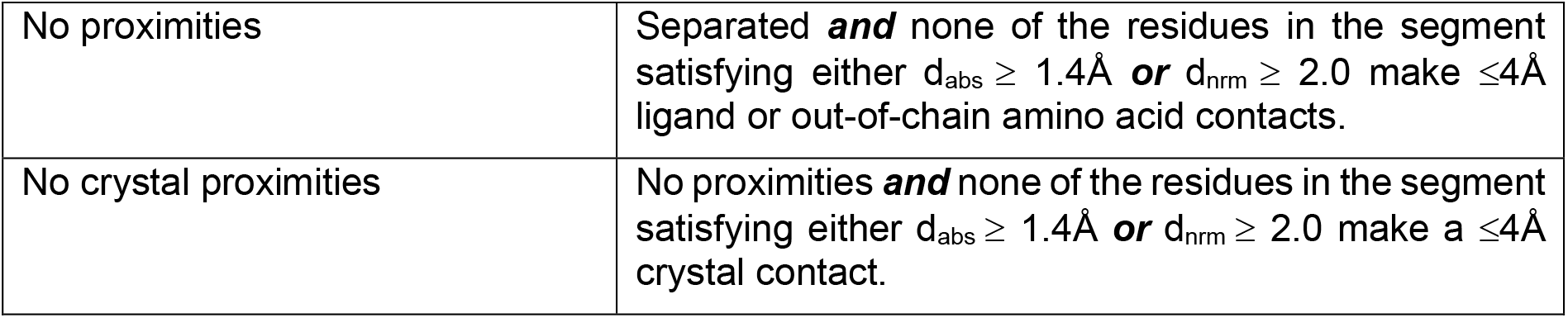

### Selection of structures for analysis in molecular dynamics simulations

Structures having separated, backbone, non-terminal altlocs, being less than 300 amino acids long in total and having no mainchain breaks were considered suitable for conventional molecular dynamics simulations. To simplify the simulations and reduce the chance of allosteric conformational changes included only structures without bound ligands. Since most models contain some small molecules from the crystallization solution, we allowed structures with any of the following ligands commonly used in crystallisation:

GOL, SO4, EDO, MSE, ACT, PEG, DMS, CL, NA, K, CA, ZN, MG.

Redundancy was removed by clustering the sequences using mmseq2 (Steinegger et al 2017) with minimum sequence identity of 0.5 and coverage 0.8 and selecting the highest-resolution structure in each unique cluster. Structures automatically selected were manually examined and further manual exclusions were made where the chain of interest was bound to a peptide or DNA/RNA or included unnatural amino acids other than selenium methionine.

### Molecular dynamics simulations

All forty-eight structures selected for molecular dynamics were prepared by created PDB files containing on the A conformation model and only the B conformation model, with all water molecules, ligands and other chains removed. To create the A conformation model all altloc B atoms, throughout the entire model were removed from the PDB and vice versa. Selenium methionine residues were mutated to methionine for recognition in the PDB2PQR server (Jurrus et al 2017) that was used to add hydrogen atoms and prepare the files for input to GROMACS.

All simulations were conducted using GROMACS software, version 2020.2 (Lindahl et al 2020). The Amber 99sb-ildnp force field (Aliev et al 2014) was applied to normal amino acids and ions, and the SPC model was applied to water molecules. After solvation in a cubic box, the addition of Cl– and Na+ ions to balance the charge, energy minimization and heating to 300K, the system was equilibrated under NVT and NPT conditions, each for 50ps. Production runs of 50ns were performed under NPT conditions, with a time step of 2fs. The temperature and pressure were maintained at 300K and 1 bar. Simulations were sampled every 20ps for each trajectory. Where the A and B conformations were found to remain stable an additional simulation, with different initial velocities, was conducted for 100ns. Glycine mutations were created in Pymol (Schrödinger and DeLano 2020) by mutating the side chain only and 50ns simulations were run using the same protocol. Heating experiments were conducted using the same protocol with the temperature set to the different values, during both equilibration and simulation stages.

### Conformation sampling

AlphaFold2 (Jumper et al., 2021) predictions were obtained by running ColabFold (Mirdita et al., 2022). AlphaFold3 (Abramson et al., 2024) predictions were obtained from alphafoldserver.com. To assess AlphaFlow-PDB, ESMFlow-PDB, AlphaFlow-MD, ESMFlow-MD (Jing et al., 2024), we sampled 50 conformations using the implementation (and weights) available at https://github.com/bjing2016/alphaflow. To evaluate Distributional Graphormer (Zheng et al., 2024), we first computed the Evoformer representations using ColabFold (Mirdita et al., 2022), and then sampled 100 conformations using the implementation available at https://github.com/microsoft/Graphormer/.

For MSA clustering + AF2 baseline, we constructed the DBSCAN-based clusters of the MSA space using the implementation provided by the authors of (Wayment-Steele et al., 2024; https://github.com/HWaymentSteele/AF_Cluster). We use the same implementation to uniformly subsample (with replacement) 10 MSA outputs with 10 or 100 sequences each (denoted as “uniform 10” and “uniform 100”).

### Visualization

Plots were created using matplotlib python package (Hunter 2007). Molecular visualization were created using Chimerax (Petterson et al. 2021).

## Data Availablity

The data from all molecular dynamics simulations will be made available on publication.

## Supplementary Figures

**Supplementary Figure 1.**
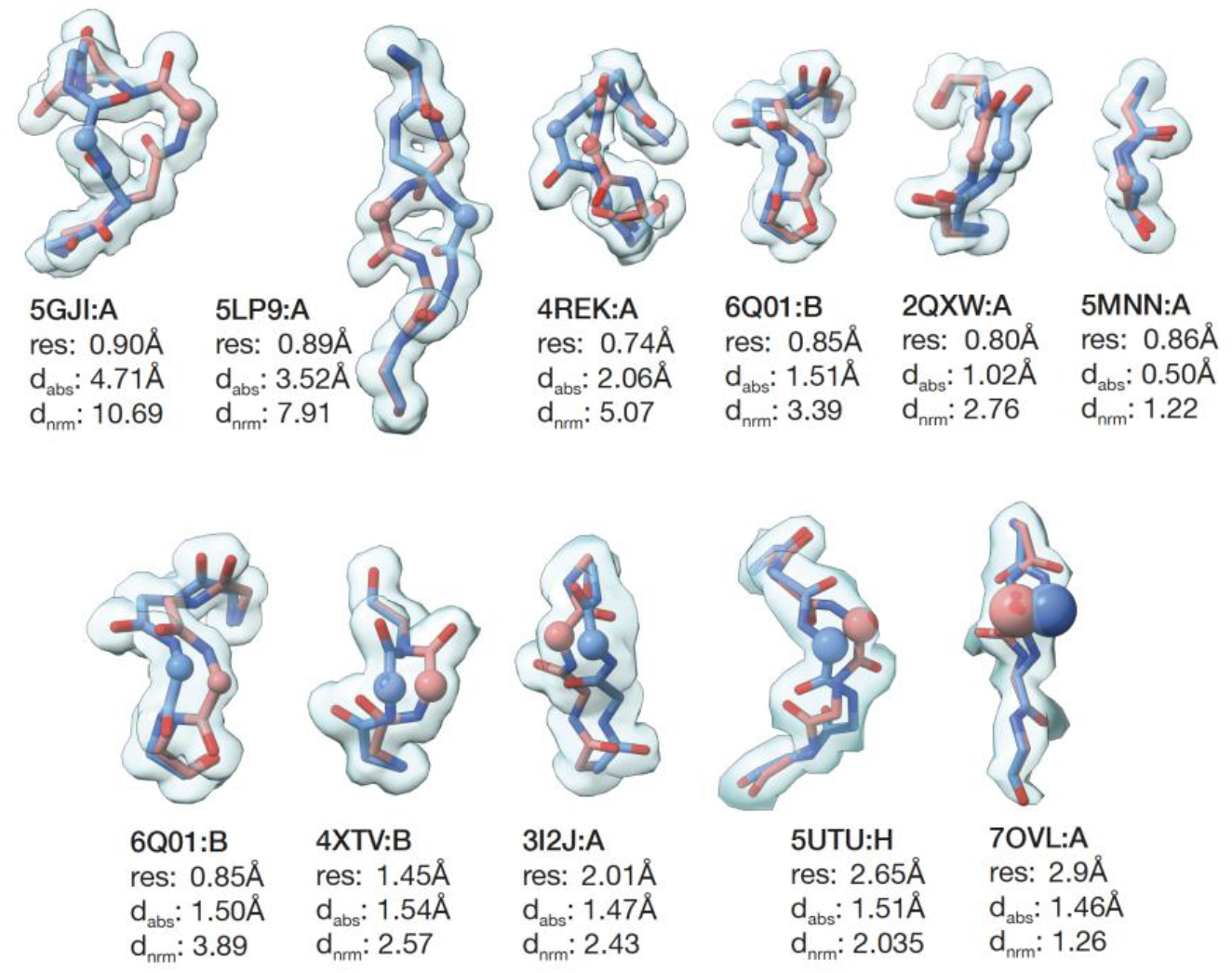
Visualization of the absolute and normalized CA-CA distances at different resolutions. First row shows varying distance at approximately constant ultra-high resolution (≤0.9Å); second row shows varying resolution at approximately constant (1.4-1.5Å absolute distance). Two modelled alternative conformations are shown as blue and red sticks. Electron densities are shown in light blue as 1σ isosurfaces. Maximally-separated (in the absolute sense) CA atoms are visualized with the radius corresponding to their B-factors.

**Supplementary Figure 2.**
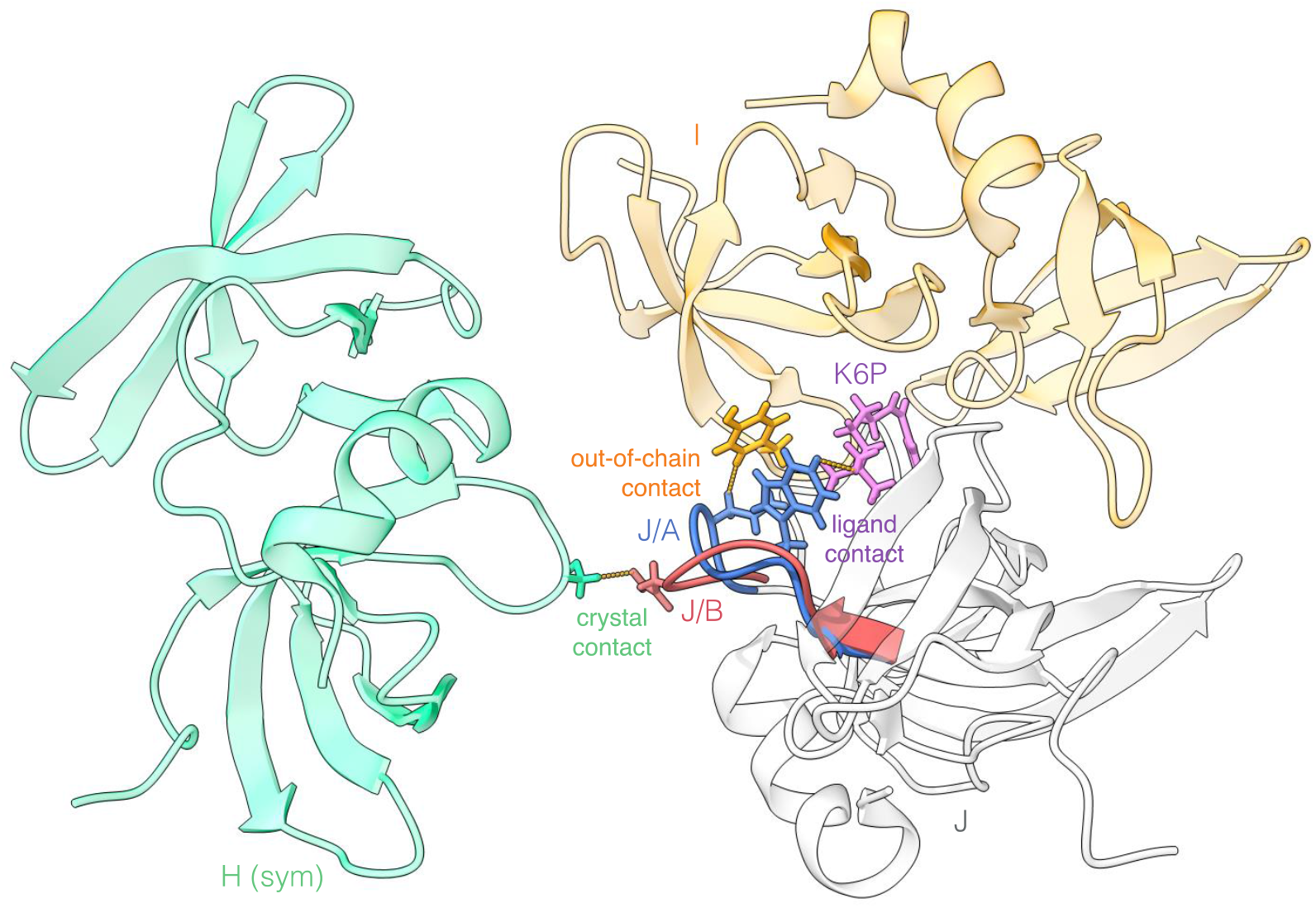
Categorisation of contact types. shown on the crystal structure 6MXX (TP53-binding protein 1 at 2.3Å resolution). Two well-separated 7 amino acid-long altloc segments in chain J (residues 1493-1499, displayed in red and blue) are in ≤0.4Å proximity of Y1523 in chain I (out-of-chain contact displayed in yellow), K6P ligand molecule (ligand contact displayed in violet), and S1497 in the symmetric replica of chain H (crystal contact displayed in green).

**Supplementary Figure 3.**
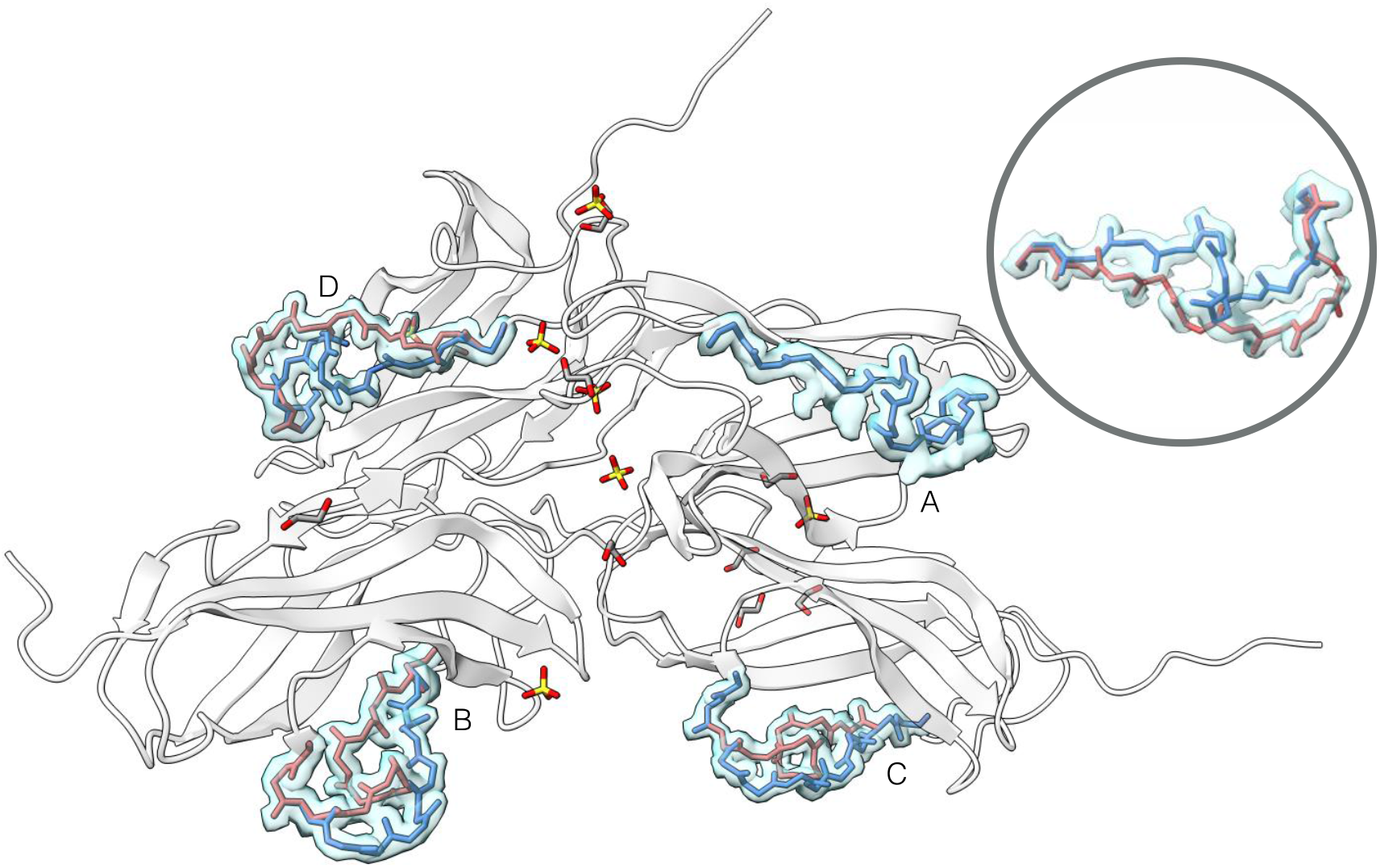
Crystal structure 4OLE. (NBR1_HUMAN protein) showing four chains with consistently modelled altloc segment of length 9 (residues 423-431) in three chains (B, C, and D). While no altloc is modelled in chain A, the counterpart from chain B fits well into the electron density (insert). 1σ density isosurfaces are shown in light blue.

